# Diel metabolic variation and the energetic demands of courtship in bioluminescent *Photeros* ostracods

**DOI:** 10.64898/2026.06.25.734310

**Authors:** Todd H. Oakley, Adriana Halvonik-Sánchez, Daniel I. Speiser, Nicholai M. Hensley

**Author notes:** **Corresponding author:** Todd H. Oakley.

## Abstract

The energetic demands of courtship are central to sexual selection, but their magnitude and temporal variation remain poorly quantified in many signalling systems. We used closed-chamber respirometry and low-light video analysis to estimate courtship-associated metabolic rates in males of the bioluminescent ostracod *Photeros* sp. “EGD.” Low-activity metabolic rate varied strongly across the diel cycle: in small vessels that constrained movement, individually measured males consumed significantly more oxygen at night than during the day. We then compared oxygen consumption across vessels that differed in opportunities for movement and courtship. Metabolic rates were highest in large vessels that permitted bioluminescent courtship displays, intermediate in medium vessels that allowed swimming but not full displays, and lowest in small vessels that constrained movement. Oxygen consumption in large vessels at night was approximately 500% of small-vessel daytime rates, 280% of small-vessel nighttime rates, and 160% of medium-vessel nighttime rates. Because measurements integrated oxygen use over multi-hour intervals, these values represent time-averaged metabolic demand rather than instantaneous costs of individual light pulses or display trains. Video analyses suggested a positive association between signalling rate and oxygen consumption, although this relationship was not statistically supported in our large-vessel dataset, which had low statistical power. Together, these results show that male *Photeros* undergo strong diel shifts in metabolic state and that the whole-animal performance required to construct bioluminescent courtship displays may often impose substantial energetic demands.

**Summary statement:** Male sea fireflies show higher nighttime metabolic rates and increased oxygen consumption during courtship-associated activities, highlighting the physiological demands of producing complex bioluminescent displays.

## Introduction

Courtship displays are among the most conspicuous and diverse products of sexual selection, reflecting variation in the signals themselves as well as the behaviours required to produce them (Byers et al., 2010; Clark, 2012). Many courtship displays require animals to coordinate movement, posture, timing, and repeated production of sensory stimuli, making courtship signalling a problem not only of communication, but also of physiological performance. The energy expenditure required for courtship displays can therefore include both production of the sensory stimulus itself and the movements that control the spatiotemporal architecture of a display. These demands may also differ across timescales: brief displays can require high instantaneous power, whereas repeated or prolonged performances can impose substantial time-averaged energetic demands (Clark, 2012; Stoddard and Salazar, 2011). Although additional experiments are required to establish fitness costs (Kotiaho, 2001), measuring metabolic rate is an important step in understanding the physiological demands of producing courtship displays. Yet the metabolic demands of most signalling systems remain incompletely quantified, especially when signal form depends on specific movements and spatial patterning.

Bioluminescent courtship displays illustrate the distinction between producing a sensory stimulus and performing the behaviour that gives the display its form. Although the energetic costs of bioluminescence remain unknown for most luminous animals, the few direct measurements available suggest that light production itself may not always dominate the energetic cost of a luminous display. In terrestrial fireflies, respirometry showed that flash production increased metabolic rate only slightly above resting levels (Woods et al., 2007). In the luminous ostracod *Photeros annecohenae*, courtship displays used only a small fraction of an individual’s stored luminescent capacity compared with defensive luminescence (Rivers and Morin, 2012). These examples leave unresolved the metabolic demands of producing spatially and temporally structured bioluminescent displays, including the swimming or flying, acceleration, stopping, secretion, and repeated pulse production that give courtship displays their form.

Luxorina, the monophyletic clade of ostracod crustaceans that use bioluminescence for courtship (Ellis et al., 2023), offers an interesting system for measuring the metabolic demands of spatiotemporally complex courtship because males generate signals using stereotyped swimming behaviours. Males in all of the dozens of Caribbean species of luxorine ostracods produce species-specific displays composed of pulses of blue light, secreted during coordinated swimming manoeuvres (Morin, 2019; Rivers and Morin, 2008). In the best-studied species, *P. annecohenae*, males initially release bright pulses while nearly stationary, then create an ascending helix of regularly spaced pulses (Rivers and Morin, 2008). Across species of Luxorina, displays vary greatly in direction, pulse duration, interpulse distance, position relative to microhabitat, and overall geometry (Cohen and Morin, 2003; Gerrish and Morin, 2016; Morin and Cohen, 2017). Males may also adjust their behaviour in relation to other males, including synchronizing or entraining with other displays (Hensley et al., 2023) and switching between signalling and sneaking tactics (Rivers and Morin, 2009). Because display geometry is generated by swimming behaviour, luxorine ostracods provide an opportunity to ask whether complex bioluminescent courtship is associated with elevated metabolic demands.

Despite published descriptions of the structure and diversity of light displays in luxorine ostracods (Gerrish, 2008; Reda et al., 2019; Rivers and Morin, 2008), no study has quantified the whole-animal metabolic demands associated with producing these courtship signals. Here, we used fibre-optic respirometry and low-light video analysis to estimate the metabolic rates associated with courtship in males of *Photeros* sp. “EGD”, an undescribed species from Panama. This species is well-suited for study because males readily produce complete courtship displays in laboratory aquaria (Hensley et al., 2023), allowing us to measure oxygen consumption under conditions that differ in the opportunity for movement and signal production. *Photeros* EGD is a close relative (Ellis et al., 2023) of the well-studied *P. annecohenae*. Although EGD differs from *P. annecohenae* in display direction and pulse structure, both species produce courtship displays by releasing trains of discrete, extracorpreal light pulses while males swim through the water column. EGD therefore provides an opportunity to extend previous work on the structure and material use of luxorine displays by quantifying the whole-animal metabolic rates associated with producing bioluminescent courtship displays. We compared metabolic rates across diel state and vessel environment, including small vessels that constrained movement, medium vessels that allowed only swimming, and larger vessels that allowed males to perform visible bioluminescent signals. We found that metabolic rates were higher at night than during the day and were highest in the large vessels in which males could perform courtship displays. Video analyses also provided preliminary evidence that oxygen consumption increased with rates of signalling activity. These results provide some of the first estimates of courtship-associated metabolic demand in bioluminescent animals.

## Methods

### Animal collection and holding

We collected *Photeros* sp. “EGD” from seagrass beds at Punta Manglar, Bocas del Toro, Panama (approx. 9.33° N, 82.25° W), in October and November of 2025. We captured some animals at night using baited PVC traps deployed in seagrass dominated by *Thalassia* and retrieved after approximately one hour. For most experimental animals, we collected males by sweep-netting through visible courtship displays at the same site (Cohen and Oakley, 2017; Hensley et al., 2023). On shore, we separated males visually and held them in containers in flow-through sea tables supplied with seawater from the field station. Ostracod colonies were fed every other day on an alternating diet of *Tubifex* worm cubes, shrimp pellets, and fish flakes. Flow-through containers were cleaned once weekly. To reduce exposure to artificial light while retaining near-natural ambient light cues, we kept animals in partially covered outdoor sea tables, using opaque tarps to block most nearby artificial lighting while allowing some exposure to the natural night sky. Animals therefore stayed at near-ambient seawater temperature and salinity until they were transferred to clean containers with 0.1 µm-filtered seawater for experiments. We allowed ostracods the opportunity to feed on a fish flake for at least 10 minutes before being transferred.

### Day–night comparison

A short pilot experiment suggested that low-activity oxygen consumption by EGD might differ between day and night. For these measurements, we used 2 mL glass vials, each containing a single male. We wrapped each vial in aluminium foil to block external light. As described in the supplemental material, the pilot experiment detected no statistical difference in oxygen consumption between wrapped and unwrapped small vessels (Figure S1). We interpret these chambers as low-activity conditions because they constrained movement and prevented courtship signalling. To test formally for diel differences in low-activity metabolism, we used a repeated-measures design in which each male was measured once overnight and once during the day in a foil-wrapped small vial (2 mL). Day and night runs were conducted back-to-back for each male, with trial order alternated across paired runs. After removing unreliable trials that we attribute to a clogged seawater filter, the analysed paired dataset included 17 males: 11 measured first during the day and then at night, and 6 measured first at night and then during the day. We estimated oxygen consumption for these matched runs using the analysis pipeline described below, and compared day and night rates using paired analyses.

### Respirometry vessels

In addition to experiments run in small vessels to determine low-activity metabolic rates, we conducted closed-chamber respirometry with medium and large vessels that differed in the space available for movement and production of courtship displays. Medium vessels were 25 mL glass scintillation vials that each housed five males. We used magnetic stir bars in the medium vessels to encourage swimming without allowing complete courtship displays. However, after two trials we observed that some animals settled on the bottom outside the influence of the stir bar. Because we lacked infrared video in the field to quantify non-luminous swimming, we discontinued medium-vessel trials after two nights and prioritized additional small-vessel replicates.

Large vessels were purpose-built by fusing graduated cylinders to the tops of screw-cap jars, both made of borosilicate glass. Large-vessel volumes ranged from 260 to 268 mL, with a height of about 22.9 cm and inside diameter about 3.8 cm. We chose this size after informal field-station trials indicated that these were approximately the smallest available vessels in which male EGD would produce their typical downward-directed courtship displays. These vessels were optically clear to permit simultaneous video recording of bioluminescent signals. Each large vessel housed 20 males because males do not produce courtship displays reliably when isolated (Rivers and Morin, 2008), and because group measurements increased the oxygen-depletion signal relative to measurement noise in the larger chambers.

### Experimental setup

We measured oxygen consumption using three multi-channel fibre-optic oxygen meters (“PyroScience bricks”; PyroScience, Aachen, Germany), including two Firesting-O2 4-channel units and one 4-channel Firesting-Pro unit, each equipped with a TDIP15 or TSUB temperature probe. Each experimental trial therefore had up to 12 oxygen channels plus three temperature channels run simultaneously. We routed fibre-optic oxygen sensors to each chamber to detect changes on OXSP5 or TROXSP5 sensor spots (PyroScience), attached to the inside of the sealed vessels. We submerged all chambers in a water bath of a single aquarium placed over a lab heater, with the water level about half the height of the large vessels. The bath temperature during experiments was maintained near ambient field conditions; approximately 27°C (range ∼26.5–28.5°C across runs, based on logged PyroScience temperature channels). For most bricks, we designated channel 1 (Ch1) as the control for microbial and background oxygen-consumption, using filtered seawater with no animals. Each brick recorded a single vessel type (small, medium, or large) during a trial, and between trials, we rotated the vessel types across bricks to minimise effects of any unquantified between-brick variation. At the end of each experimental trial, we dried the ostracods at 60°C for 24–48 h. Later, we estimated the dry mass (M) of individuals from the small chambers with a Mettler Toledo UMX2 ultra micro-balance (Mettler Toledo LLC, Columbus, USA) and estimated bulk masses of individuals from the medium and large chambers with a Mettler Toledo XS104 analytical balance.

We logged raw oxygen traces (µmol O₂ L⁻¹ versus time) and temperature from each channel at regular intervals (2 or 5 s) using PyroScience acquisition software. We collected data using the ‘respirometry’ package in R (Birk, 2025). Subsequent processing used a custom analysis pipeline implemented in R (consumption_rate.R) that was called from a Jupyter notebook (“B_batch_respirometry”) via Python. Briefly, for each “run” (defined as a brick × trial combination), we supplied the script with salinity (33, in practical salinity units from nearby monitoring station readings), identities of the channel used as a control with no animals (usually Ch1), animal channels (typically Ch2–Ch4), chamber volumes, number of animals in the vessel and measured M of test animals per channel, and the time window to estimate rates (generally 1–8 h after sealing). The R script fit linear regressions to oxygen concentration versus time for each channel over the specified window. Microbial/background respiration from the control channel was converted to the same units using the control volume and subtracted from each animal channel recorded with the same brick. We expressed oxygen consumption as mass-specific metabolic rate (µL O₂ mg⁻¹ h⁻¹) and applied a quarter-power mass correction using dry mass (M), following standard allometric scaling approaches (Glazier, 2005; Savage et al., 2004). In this correction, M is dry body mass in grams and the scaling exponent was set to −0.25. The Python notebook aggregated per-channel summaries across runs into a single long-format table containing uL_mg_hr, temp_C and metadata (e.g. trial ID, vessel type, date, day/night treatment) for downstream statistical analyses. Code and data for all analyses are available at https://github.com/ucsb-oakley-lab/signal_respirometry.

### Video analysis of signalling activity rate

To quantify bioluminescent courtship activity (a parameter we call *signal_rate*) inside the large vessels, we recorded each trial using a GoPro HERO13 Black action camera (GoPro Inc., San Mateo, CA, USA) mounted outside the water bath and aimed laterally at the capped graduated cylinders. We set cameras to “Star Trails” mode, which uses long-exposure frame accumulation, with all automatic settings otherwise left at manufacturer defaults (0.03 fps). For each trial, recording began when we started metabolic measurements and extended through the entire trial, with the exception of trial 1, when the camera battery was exhausted after about 2.5 hours. We used wall power after the first night. Later, we analysed videos using custom Python scripts built with OpenCV (available on GitHub). In brief, a script first converted frames to greyscale, before applying a background-subtraction and thresholding routine to isolate bioluminescent events on each frame. In particular, we ignored the orange light from the fibre-optics, which we also assume did not affect behaviour of the animals. We then used contour (e.g. contiguous regions of bright pixels) detection to identify discrete luminous “pulses” in each frame, with each detected contour assumed to be a single bioluminescent pulse. We then identified the locations of multiple large vessels in each video to allow separate estimates of signalling rate for each vessel, and extracted a time series of pulse counts per fixed time bin (Figure S2), which we used to quantify *signal_rate* as our estimate of signalling activity. We then matched these estimates of signalling activity to oxygen-consumption estimates for each large vessel to attempt to relate signalling propensity to metabolic rate.

### Comparative context

To place the metabolic rates associated with courtship of *Photeros* sp. EGD in a broader context, we replotted comparative data from a variety of animals (not including ostracods) on resting and signalling metabolic rates compiled by Stoddard and Salazar (2011) and added two *Photeros* points from the present study. We also updated this comparative dataset with post-2011 studies identified through a targeted literature search, followed by manual verification of original papers, supplementary materials and public data repositories. Searches combined terms for metabolic rate or respirometry with animal signalling terms, including calling, song, whistle, click, echolocation, hissing, stridulation, and vibrational signalling. We retained candidate studies only when signalling and baseline metabolic rates could be expressed as mass-specific oxygen consumption in the same units as the original dataset, ml O₂ h⁻¹ g⁻¹. Both *Photeros* points used the mean metabolic rate measured in large-vessel trials as the courtship-associated rate, but differed in the low-activity baseline used for comparison: daytime small-vessel metabolism and nighttime small-vessel metabolism. We included both of these baselines because *Photeros* metabolic rate differed strongly across the diel cycle, making daytime resting metabolism and nocturnal low-activity metabolism biologically distinct reference points. We treated this cross-species comparison as descriptive rather than inferential because the literature values are phylogenetically clustered and differ across original studies in methods, signal modality, baseline definition, and the degree to which signalling rates represent instantaneous signal production versus time-averaged behaviour.

## Results

### Low-activitymetabolic rate increased at night

Individual males measured in small, foil-wrapped chambers showed a consistent diel difference in oxygen consumption. Across paired day–night measurements of the same individuals, metabolic rate was higher at night than during the day (Fig. 1). This pattern is significant using both non-parametric and parametric paired tests: nighttime rates exceeded daytime rates in most paired comparisons, and the difference was significant using a Wilcoxon signed-rank test (W = 5.000, p = 0.0002) and a paired t-test (t = 5.053, p = 0.0001). Because water temperature varied slightly among trials, we tested whether temperature explained the paired day–night difference in metabolic rates of the small vessels. In a random-intercept mixed model, the effect of night remained strongly positive after accounting for temperature (Figure S3). Thus, the difference in low-activity metabolism between day and night was not explained by the small temperature variation among paired small-vessel trials. Additionally, because animals were confined individually in the smallest vessels and shielded from external light, we interpret these values as low-activity metabolic rates, without courtship or much active swimming. The magnitude of this diel shift indicates that baseline metabolic state in *Photeros* differs between day and night, even when animals are in the dark and not able to signal. This result is important for interpreting courtship costs because daytime resting metabolism alone could underestimate the relevant nocturnal baseline for animals that naturally display at night.

**Figure 1.**
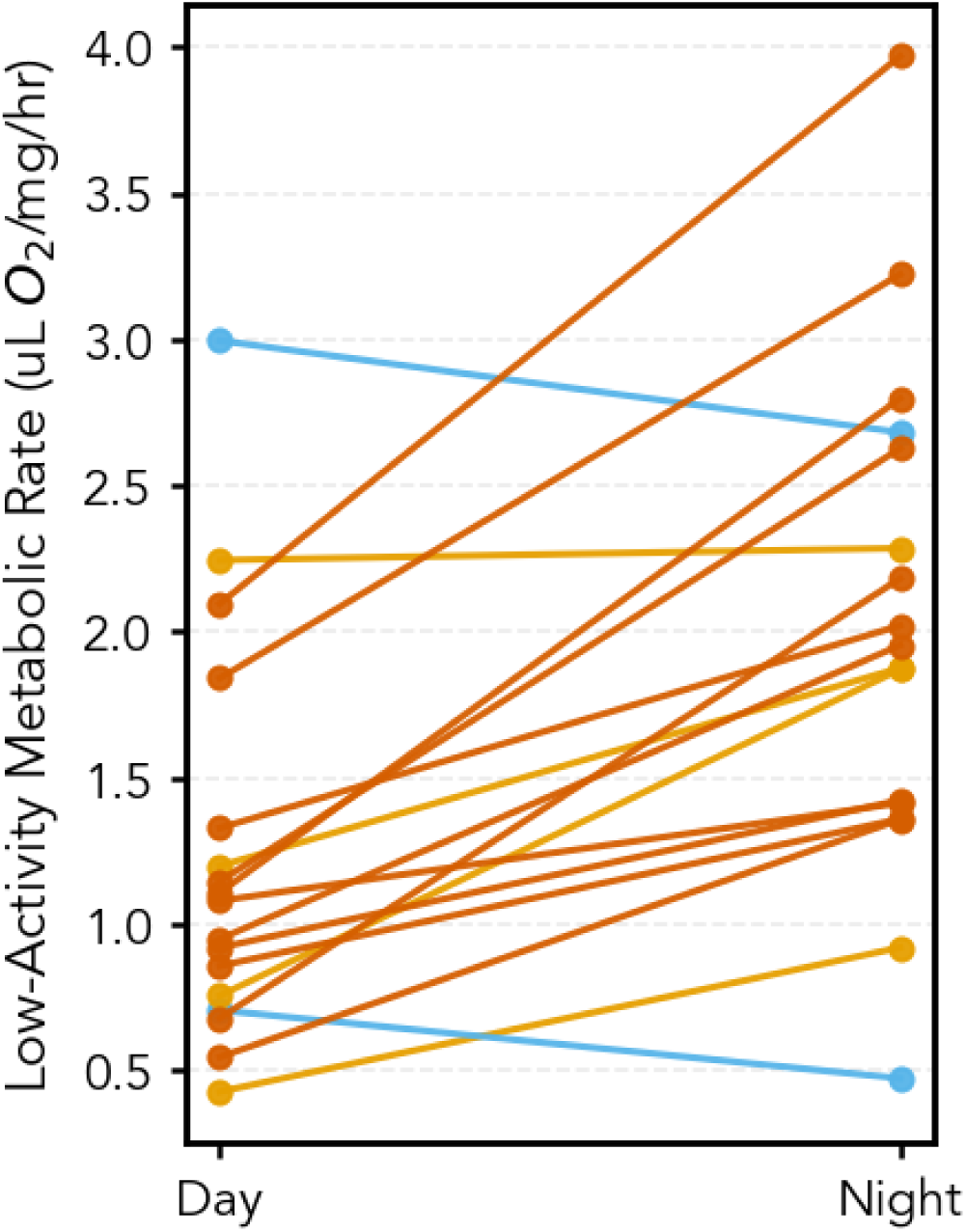
Paired day-night metabolic rates for the same *Photeros* individuals measured across sequential trial pairs. Each line connects day (left) and night (right) measurements of a single male in a small vessel covered in foil. Orange lines indicate pairs in which metabolic rate was higher at night than during the day; blue lines indicate pairs in which metabolic rate was equal to or higher during the day than at night. Shade intensity indicates trial order: light shades represent pairs that started with day measurements and dark shades represent pairs that started with night measurements. Across all filtered paired comparisons (n = 17; 11 pairs measured day first and 6 pairs measured night first), nighttime metabolic rate was significantly higher than daytime metabolic rate (Wilcoxon signed-rank: W = 5.000, p = 0.0002; paired t-test: t = 5.053, p = 0.0001). Notebook with all code at https://github.com/ucsb-oakley-lab/signal_respirometry/blob/main/notebooks/Figure1_SlopeGraph.ipynb

### Metabolic rates increased with vessel size and opportunity for courtship

Mass-specific metabolic rates varied among chamber treatments (Figure 2). Mean metabolic rate was highest in large-vessel trials at night (5.84 ± 2.44 SD µL O₂ mg⁻¹ h⁻¹; n = 8), intermediate in medium vessel trials at night (3.61 ± 1.02; n = 6), and lower in small vessels measured at night (2.10 ± 1.08; n = 26) or during the day (1.18 ± 0.70; n = 18). Rates in the large vessels were therefore 1.6-fold higher than rates in medium vessels, 2.8-fold higher than rates in small vessels at night, and 5.0-fold higher than small vessels during the day. We fit an ordinary least-squares model and used HC3 heteroskedasticity-robust standard errors to reduce sensitivity to unequal residual variance, using the trials in large vessels as the reference group and testing directional hypotheses that metabolic rates in large vessels exceeded those of each comparison group. Metabolic rates of animals in large vessels were significantly higher than rates of animals in medium vessels at night (difference = 2.23 µL O₂ mg⁻¹ h⁻¹; one-tailed p = 0.015), rates in small vessels at night (difference = 3.75 µL O₂ mg⁻¹ h⁻¹; one-tailed p = 3.82 × 10⁻⁵), and small vessels during the day (difference = 4.66 µL O₂ mg⁻¹ h⁻¹; one-tailed p = 3.26 × 10⁻⁷). To control the family-wise error rate (mulitple comparisons) across the three directional tests, we applied the Holm correction; all contrasts remained significant (Holm-adjusted p = 0.015, 7.63 × 10⁻⁵, and 9.79 × 10⁻⁷).

**Figure 2.**
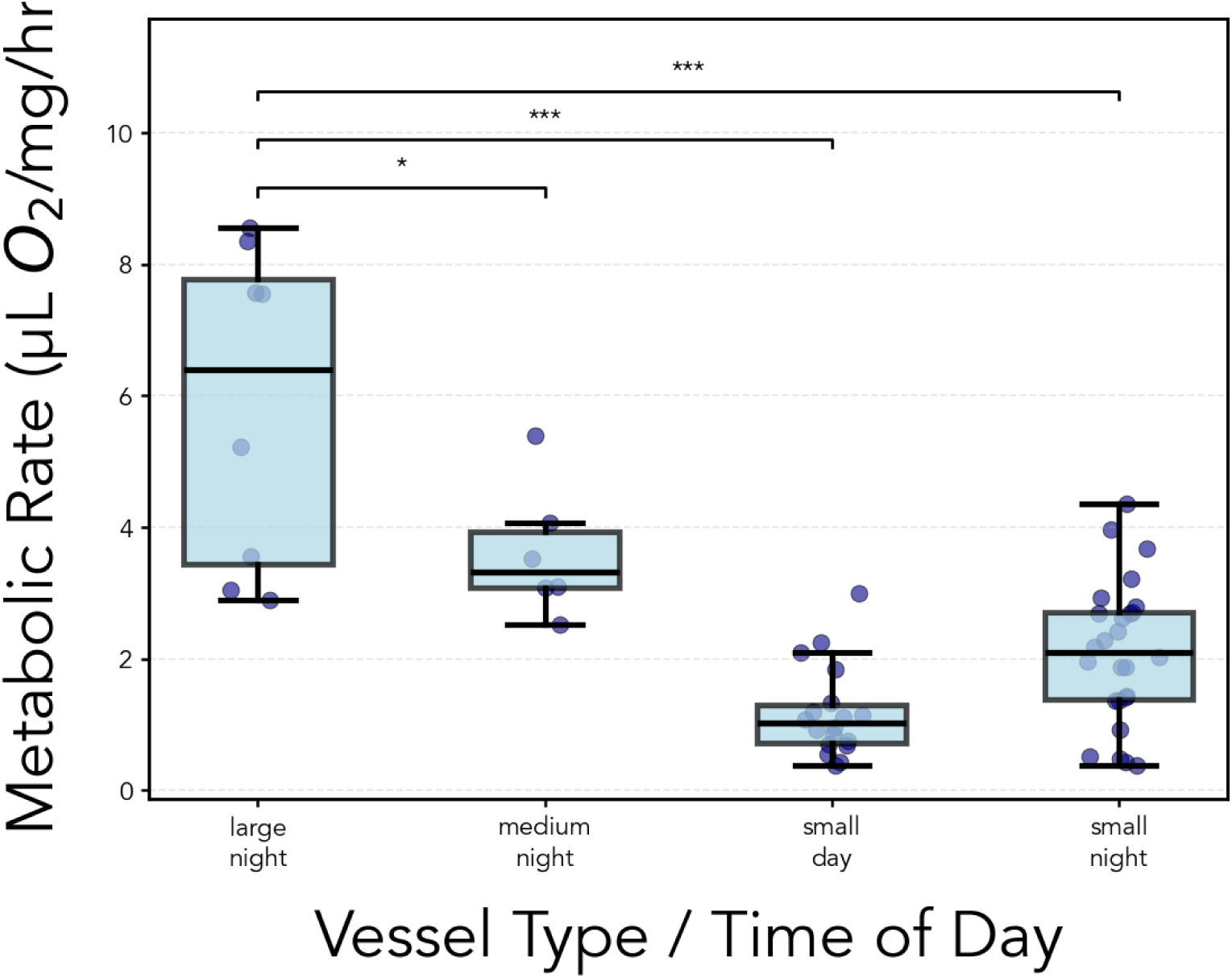
Mass-specific metabolic rate (µL O₂ mg⁻¹ h⁻¹) of *Photeros* EGD measured across vessel type and time of day. Metabolic rates were highest in large vessels at night, intermediate in medium vessels at night, and lowest in small vessels; within small vessels, rates were higher at night than during the day. We interpret these differences under the assumption that activity varied by treatment: courtship signalling in large-night trials, swimming without visible signalling in medium-night trials, and reduced movement in small vessels. Boxes show medians and interquartile ranges; whiskers show the non-outlier range; points show individual observations. Sample sizes were: large vessels, n = 8 with 20 animals per vessel; medium vessels, n = 6 with 5 animals per vessel; small-day, n = 18; small-night, n = 26, with one male per small vessel. Metabolic rates of animals in large vessels at night exceeded the rates of all other groups in directional contrasts from an ordinary least-squares model with HC3 heteroskedasticity-robust standard errors, using large-night as the reference group. Holm-adjusted one-tailed p-values were 0.015 for large-night versus medium-night, 7.63 × 10⁻⁵ for large-night versus small-night, and 9.79 × 10⁻⁷ for large-night versus small-day. Note the illustrated data from small vessels include the paired trials illustrated in Figure 1 plus additional unpaired, mainly nighttime trials.

For sensitivity analyses, we repeated the same comparisons after data points we removed due to the negative estimates of oxygen consumption in large vessels, which we attribute to high background respiration in control chambers caused by insufficiently filtered seawater. In this full-dataset analysis, the comparison between large vessels and medium vessels became non-significant, but oxygen consumption rates in large vessels remained significantly higher than consumption rates of animals in both small-vessel groups (Figure S5). We also repeated statistical comparisons from Figure 2 with temperature included as a covariate. Comparisons using temperature remained qualitatively unchanged and remained significant, suggesting that the main pattern of differences in metabolic rate among vessel sizes was not explained by the small temperature variation among trials (Table S1). Overall, we interpret differences in consumption rates to be mainly related to behavioural opportunities provided by each vessel type. Small vessels constrained movement, allowing estimates of low-activity routine metabolism, medium vessels allowed some nocturnal swimming but not courtship signalling, and large vessels allowed males to perform visible courtship displays. The elevated metabolic rates in large vessels therefore indicate that courtship-associated behaviour carries substantial metabolic demands relative to both low-activity and active baselines.

### Higher rates of oxygen consumption in larger vessels that allowed courtship

The magnitude of the increase of metabolic rates in the largest vessels depends on the baseline used for comparison. Metabolism in large vessels was approximately 495% of the daytime low-activity baseline and approximately 279% of the nocturnal low-activity baseline. Compared to medium sized vessels, which provide the closest available comparison for nocturnal activity without reliable courtship signalling, metabolic rates in large vessels was approximately 162% of this active nocturnal baseline. These comparisons indicate that the energetic demand of courtship-associated behaviour is substantial, even under the more conservative comparison to active nocturnal animals. In addition, because we measured oxygen consumption over multi-hour intervals, metabolic rates in large vessels represent time-averaged courtship-associated metabolism rather than the instantaneous cost of individual pulses or individual display trains. The measured values include periods of signalling, swimming, non-signalling activity, and recovery (Rivers and Morin, 2008; Rivers and Morin, 2009).

### Signalling activity was positively associated with oxygen consumption but limited by sample size

Video analysis of large-vessel trials allowed us to estimate bioluminescent signalling activity (*signal_rate*) and relate it to oxygen consumption. Across measurements from individual large vessels, oxygen consumption tended to increase with estimated signalling rate, but the filtered dataset did not provide strong statistical support for this relationship. Regression showed a positive association between signalling rate and metabolic rate (R² = 0.447), but the relationship was not significant at α = 0.05 (p = 0.070; SE = 1.085). For a number of reasons, this result should be interpreted cautiously. First, temperature covaried strongly with signalling rate in the large-vessel trials (r = 0.873), and accounting for temperature made the apparent positive association between *signal_rate* and metabolic rate ambiguous (Figure S4). This covariation could be important because crustacean respiration is generally temperature-sensitive (Ivleva, 1980), and recent work found a significant positive effect of temperature on routine metabolic rate in two swimming podocopid ostracods (Cruz-Vila et al., 2026). Even though the paired small-vessel analysis above indicated that temperature differences among trials did not explain the diel increase in low-activity metabolism, we do recognize the confound of temperature and *signal_rate* as an important limitation of the regression analysis comparing metabolic rate and signaling rate. Second, we also excluded some measurements of metabolic rates (Figure S6) because they produced negative estimates of oxygen consumption, which we attribute to high background respiration in control chambers caused by insufficiently filtered seawater. One night, our filter was clogged and we attempted experiments with unfiltered water. Removing these trials reduced the sample size and therefore the power of the analysis. Third, courtship activity was uneven across nights, with the highest overall signalling activity occurring on the first night of trials when temperatures were highest. A power analysis indicated that approximately 12 additional replicate measurements would be needed to detect the observed effect at p < 0.05 under current assumptions. In the end, we consider the filtered dataset to be consistent with the hypothesis that more signalling increases oxygen consumption, but additional trials with low background respiration, more evenly distributed signalling activity, and tighter temperature control will better test the hypothesis and estimate the slope precisely.

### Courtship metabolism in comparative context

We placed the metabolic cost of signalling in *Photeros* in a broader comparative context by adding our estimates to published measurements of resting (which we term low-activity) and signal-associated metabolic rates across animal communication systems (Figure 4). We replotted the Stoddard and Salazar dataset and added post-2011 studies identified through our literature update, including additional measurements from katydids, dolphins, bats, leafhoppers, and vipers. These updated points broadened the comparison to include whistling, echolocation clicks, vibrational advertisement calls, and defensive hissing. The position of *Photeros* in the plot depended on which estimate of low-activity metabolic rate that we considered to be the baseline. Using daytime low-activity metabolism placed courtship-associated metabolism at approximately 5-fold resting levels, whereas using nocturnal low-activity metabolism placed courtship-associated metabolism at approximately 2.8-fold resting levels. Thus, even without accounting for the fraction of time males spent actively signalling, courtship-associated metabolism in *Photeros* fell within the range of elevated metabolic rates reported for other energetically demanding displays. The nocturnal baseline comparison is likely the more behaviourally relevant reference because courtship occurs at night, when low-activity metabolism was already elevated.

**Figure 3.**
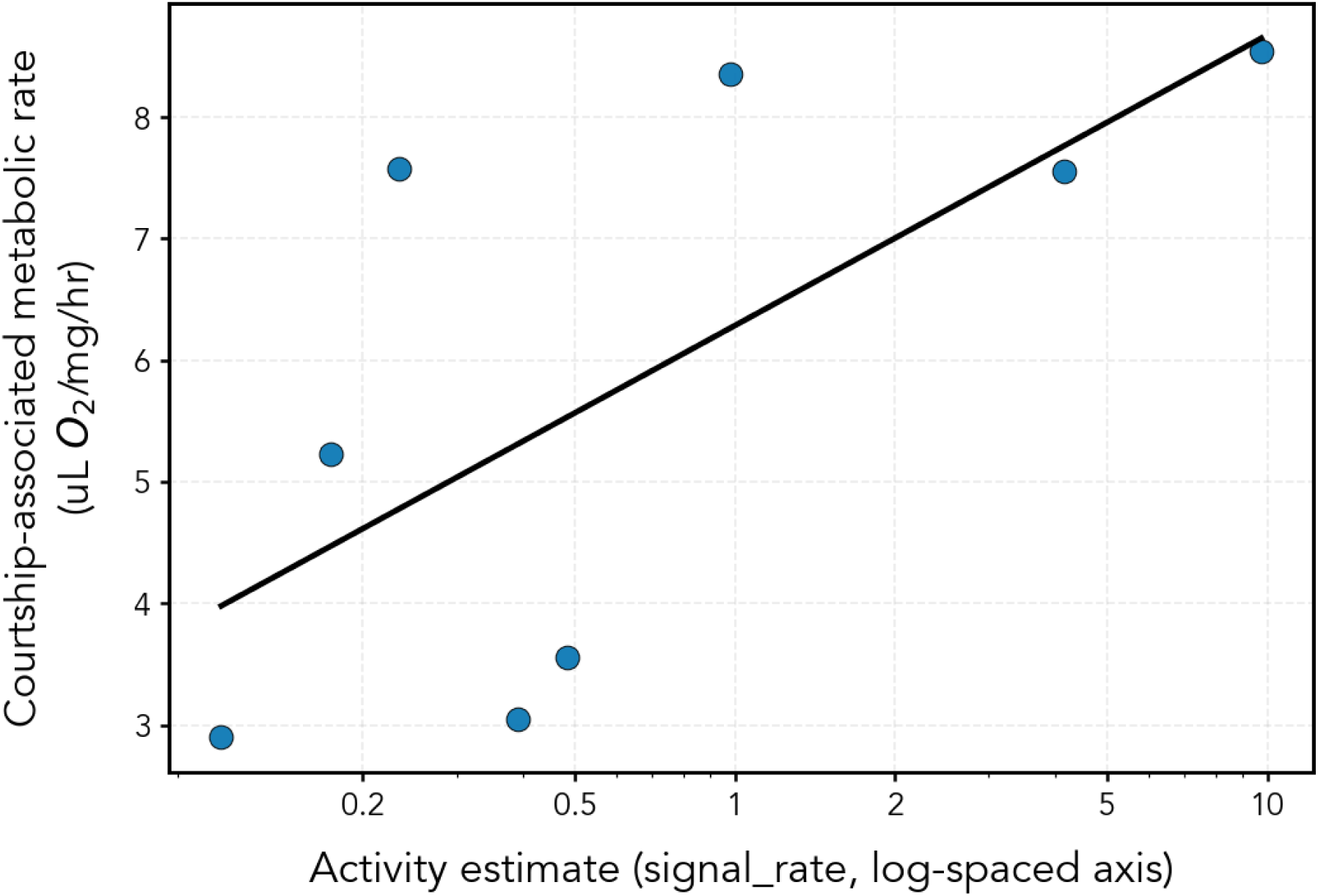
Relationship between estimates of signalling activity and metabolic rate in large-vessel trials. Points represent individual trials, with routine metabolic rate (RMR) plotted against activity estimate (signal_rate = mean contours (small contiguous regions of bright blue pixels) detected on video per 60s; log-scaled x-axis). The solid line shows the least-squares fit to *CAMR*∼log10 (signal_rate). The relationship is positive (slope = 2.39), with moderate explained variance (*R*^2^ = 0.45) and marginal statistical support (*p* = 0.070, *n* = 8). Post hoc power analysis (Fisher-z approximation) indicated 0.44 power to detect the observed association; approximately n = 16 (8 additional) and n = 20 (12 additional) would be needed for 0.80 and 0.90 power, respectively.

**Figure 4.**
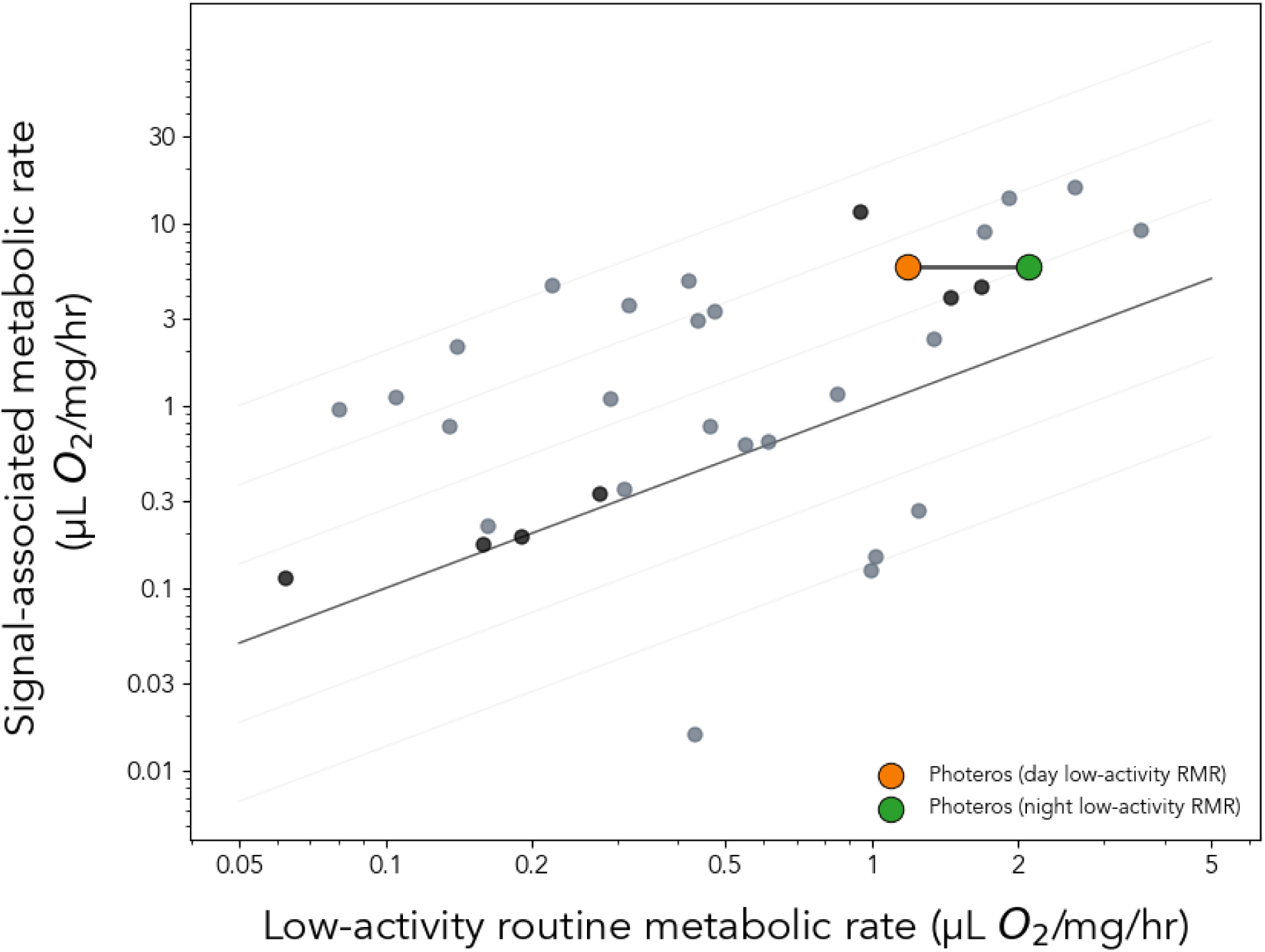
Published estimates of resting and signalling metabolic rates across animal communication systems are replotted from (Stoddard and Salazar, 2011), with additional post-2011 measurements from the literature (black data points) and two *Photeros* sp. “EGD” points added for comparison. The two *Photeros* points use the same large-night metabolic rate as the courtship-associated value, but differ in the baseline used for resting metabolism: daytime low-activity metabolism and nighttime low-activity metabolism. Diagonal reference lines indicate equivalent proportional increases above rest. *Photeros* values are time-averaged across multi-hour trials rather than instantaneous pulse costs (see Results).The comparison is therefore intended to provide descriptive context rather than a formal phylogenetic meta-analysis.

### Summary of results

Together, these analyses show that metabolic rate of *Photeros* varies across diel and behavioural contexts. First, the low-activity metabolic rate was higher at night than during the day. Second, metabolic rates increased across vessel treatments in a pattern consistent with greater behavioural opportunity and courtship activity. Third, trials in courtship-associated large vessels produced the highest metabolic rates, exceeding both resting and active nocturnal baselines. Finally, signalling activity showed a positive but underpowered association with oxygen consumption. These results support the conclusion that bioluminescent courtship in *Photeros* imposes substantial metabolic demands, most likely because light production occurs together with complex movement patterns, rather than because biochemical light production alone has a substantial metabolic demand.

## Discussion

Bioluminescent courtship displays are visually conspicuous, but their energetic significance may lie not only in light production itself, but also in the whole-animal performances required to construct signals in space and time. In many luminous species, signalers do not simply emit light; they swim or fly, respond to social cues, and place luminous pulses or flashes into intricately structured displays (Morin, 2019; Oakley, 2025; Stanger-Hall and Oakley, 2019). Our results therefore extend previous work on the cost of light production itself (Rivers and Morin, 2012; Woods et al., 2007) by emphasizing the additional metabolic demands of behaviours associated with bioluminescent courtship. Viewed this way, the elevated oxygen consumption measured in signalling conditions is best interpreted as courtship-associated metabolic demand: an integrated outcome of nocturnal physiological state, locomotion, social interactions, secretion, and light production.

This interpretation is supported by two features of the data. First, male *Photeros* had higher metabolic rates at night than during the day when confined in small darkened vessels. Second, metabolic rates were highest in large vessels that allowed visible courtship displays compared to both intermediate vessels that allowed swimming but not full displays, and the smallest vessels that restricted movement the most. Together, these patterns indicate that courtship-associated metabolism in *Photeros* EGD reflects both a nocturnal shift in physiological state and the additional energetic demands of movement and display production. This interpretation is also consistent with the structure of courtship displays by luxorine ostracods. In these ostracods, males do not simply release light; they construct species-specific spatial and temporal patterns by swimming and secreting discrete luminous pulses. In a related species, *P. annecohenae*, males produce an initial stationary phase of pulses followed by a helical phase in which they place pulses at nearly regular spatial and temporal intervals as males swim rapidly upward (Rivers and Morin, 2008). In *Photeros* sp. “EGD,” males produce downward displays, often in collective waves, with displays also influenced by environmental and social cues (Hensley et al., 2023). In all Luxorina species studied to date, the luminous signal is inseparable from movement: the geometry of the display is generated by the animal’s trajectory through the water.

Although here we emphasize movement and whole-animal performance, light production and secretion are still part of the energetic accounting. Cypridinid bioluminescence also involves oxidation of secreted luciferin, and the visible light output of each pulse depends on the amount of substrate oxidized, the activity of luciferase, and the quantum yield of the reaction. In addition, luciferin, luciferase, and associated secretory products must be synthesised and stored before signalling, so some energetic costs of courtship are paid before the display itself. Our measurements cannot partition oxygen consumption among these components. We did not measure absolute photon output, quantum yield, substrate depletion, or oxyluciferin production, and our respirometry integrates oxygen consumption across groups of active males over hours. Thus, light production almost certainly contributes to the metabolic rates measured in the large vessels, but the present data cannot partition that demand among biochemical light production, swimming, acceleration, secretion, and other courtship-associated behaviours.

Our video analyses suggest a link between signalling activity and oxygen consumption, but this result should be interpreted cautiously. Across large vessels with different rates of signalling, oxygen consumption tended to increase with estimated signalling rate, but the relationship was not statistically significant, especially when accounting for temperature differences. Several features of the experiment limit inference: the overall number of large-vessel observations was small, some trials were excluded because high background oxygen consumption in control chambers produced unreliable estimates, and signalling activity was high mainly during the first night of trials, perhaps because of lunar phase or animal condition. Even so, the direction of the association is consistent with the expectation that groups producing more displays should have higher metabolic rates. More importantly, the combined respirometry and video approach developed here provides a framework for future studies that connect metabolic physiology to display variation. With larger datasets, this approach could also test whether metabolic rate increases approximately linearly with display production or instead approaches a performance ceiling at high signalling rates. Because EGD males participate in social signalling aggregations and can adjust display behaviour in relation to nearby signals (Hensley et al., 2023), metabolic capacity may influence not only whether males signal, but also how often they display, how many pulses they produce, how fast they swim, and how often they switch among signalling and non-signalling tactics.

Comparative work will be needed to determine whether the metabolic costs of courtship displays in Luxorina are relatively high or low compared to those of the movement-based mating behaviours of other animals. Terrestrial fireflies also combine bioluminescent signalling with locomotion, but ostracods are smaller, millimetre-scale aquatic crustaceans construct signals in a very different physical environment. Displaying ostracods operate at intermediate Reynolds numbers and males may slow or stop during pulse secretion to maintain discrete luminescent packets rather than leaving streaking light trails behind the swimmer (Rivers and Morin, 2008). This physical context also highlights the distinction between time-averaged energy expenditure and instantaneous metabolic power (Clark, 2012). Our oxygen-consumption estimates were averaged over multi-hour trials, whereas males likely spent only a fraction of that time producing visible display trains. A duty-cycle-corrected estimate would require estimating the fraction of each trial spent actively displaying and using that time budget to infer the metabolic power associated with active display periods. Because our sample sizes were low and our temporal measurements were imprecise, we did not attempt this correction. Instead, our estimates should be interpreted as trial-averaged courtship-associated metabolic rates. If visible display trains occupied only a small fraction of each trial, then the instantaneous metabolic demand of active displays could be higher than the time-averaged values reported here. Thus, although bioluminescence is the visually distinctive feature of luxorine displays, the locomotor behaviours that position, separate, and spatially organise pulses may be equally or more important contributors to metabolic demand. Interestingly, complex swimming behaviours may have preceded the evolution of luminous courtship because some non-luminous myodocopid ostracods perform conspicuous courtship movements, including helical swimming (Arenz et al., 2018; Lum et al., 2008), suggesting that the motor components of display behaviour may be evolutionarily older than the use of light as a sexual signal.

Previously published measurements of ostracod metabolism provide useful taxonomic context, but they should not be treated as direct comparisons to our courtship-associated measurements. Previous studies measured oxygen consumption in a range of marine and freshwater ostracods, including planktonic halocyprids (Ikeda, 1990; Kaeriyama and Ikeda, 2004), podocopids (Cruz-Vila et al., 2026), and cypridinids (Donnelly et al., 2004), across broad differences in temperature, body size, activity state, and mass correction. These values suggest that the rates we measured for *Photeros* fall within the broader physiological range reported for ostracods, with courtship-associated values near the upper end of many reported routine or active metabolic rates. However, previous ostracod studies generally measured resting, routine, swimming, or environmentally induced metabolic rates rather than metabolism during male courtship displays in a social signaling context. The strongest comparison in the present study is therefore internal: males of the same *Photeros* species consumed more oxygen at night than during the day, and oxygen consumption was highest in vessels that allowed bioluminescent courtship displays.

The diel shift in low-activity metabolic rate we observed in EGD is biologically important in its own right. Among crustaceans, diel patterns in metabolism vary across species, physiological traits, and environmental contexts. For example, oxygen consumption and ammonium excretion showed no significant diel pattern across the migration cycle in the vertically migrating copepod *Pleuromamma xiphias*, even though fecal pellet production peaked at night and metabolic enzyme activities varied with time of day (Tarrant et al., 2021). By contrast, diel or circadian patterns in oxygen consumption and metabolic physiology exist in other planktonic crustaceans, including Antarctic krill and the copepod *Calanus finmarchicus* (Häfker et al., 2017; Teschke et al., 2011). Diel activity rhythms also are present in myodocopid ostracods: *Cylindroleberis mariae* swims actively in the water column at night but rests during the day in self-made nests, where individuals experience confined, hypoxic conditions that may contribute to metabolic depression or torpor-like behaviour (Corbari et al., 2005). In *Photeros* EGD, we found higher nighttime oxygen consumption to persist in small, darkened vessels that strongly constrained movement and prevented courtship signalling. This result suggests that males enter a higher-metabolism nocturnal state, even before the additional demands of swimming and signalling. Anecdotally, luxorine ostracods handled during the day often appear less responsive to disturbance than those handled at night, suggesting that daytime metabolic rates may reflect a quiescent or sleep-like state rather than simply inactivity imposed by the darkened chamber. Formal tests of diel responsiveness, arousal thresholds, spontaneous movement, and endogenous clocks would help determine whether daytime metabolic depression represents a regulated behavioural state associated with nocturnal courtship.

Several limitations are important for interpreting these results. First, as noted above, fibre-optic oxygen measurements are well suited for estimating integrated oxygen consumption across a trial, but are unlikely to provide the temporal resolution needed to estimate rapid changes in power during individual courtship manoeuvres. Second, vessel treatments differed in multiple ways, including not only space for movement and therefore opportunity to signal, but also group size, and social context. We therefore interpret the large-vessel treatment as courtship-associated rather than as an isolated manipulation of signalling. Third, our video analysis quantified luminous pulses but not invisible swimming, so silently moving males, including potential sneakers, would contribute to oxygen consumption without being counted in the signal rate parameter. Future experiments could combine infrared and low-light video to count both visible display trains and non-luminous participants in courtship swarms, estimate apparent swimming speeds and movement paths, and relate these behavioural measures to time-averaged metabolic rates.

Together, these results show that bioluminescent courtship in *Photeros* is associated with elevated whole-animal metabolic demand. Our measurements pool oxygen consumption from light production, secretion, movement, and social courtship behaviour, and therefore estimate courtship-associated metabolism rather than the cost of bioluminescence alone. In luxorine ostracods, light is placed into the environment by swimming animals, and the display’s temporal and spatial structure depends on movement. If individual males differ in their capacity to sustain elevated metabolism associated with courtship, these physiological differences could contribute to variation in display rate, pulse number, swimming performance, or mating tactic use. Such variation could make these display traits candidates for honest or performance-based signals, although testing this possibility will require linking individual metabolic capacity to display traits, female responses, and mating success (Byers et al., 2010; Maynard-Smith and Harper, 2003). Bioluminescence makes luxorine courtship performances conspicuous, but their physiological demands extend beyond the cost of light itself to include enzyme and substrate production, secretion, movement, and social interaction. Luxorine ostracods therefore offer a useful system for studying how metabolic physiology, locomotor performance, and social signalling interact during the evolution of complex courtship displays.

## Acknowledgements

We thank Dr. R. Collin and P. Gondola at the Smithsonian Tropical Research Institute for support, and we are grateful to members of the STRI Bocas del Toro Research Station for logistical assistance during fieldwork. Thanks to C. Guerra for crafting the custom glass vessels. We also thank C. McKinley for helpful discussions during the planning of this work. Animals were collected with permission from the Panamanian government (MiAmbiente permit # ARB-0127-2022).

## Competing interests

The authors declare no competing interests.

## Funding

This work was supported by the Human Frontier Science Program [RGP021/2024-101 to T.H.O. and RGP016/2023 to D.I.S.]; and the University of California, Santa Barbara Faculty Research Grants Program [to T.H.O.]. N.M.H. was supported by a Herchel-Smith Postdoctoral Fellowship from the University of Cambridge.

## Data and resource availability

All data and code generated for this study will be made publicly available at the time of publication in a GitHub repository: https://github.com/ucsb-oakley-lab/signal_respirometry. Upon publication, an archived version of the repository will be deposited in Zenodo and cited with a DOI in the reference list: [Zenodo DOI].

## Notes

https://github.com/ucsb-oakley-lab/signal_respirometry

### Competing Interest Statement

The authors have declared no competing interest.

