## Supplemental Information for "Diel metabolic variation and the energetic demands of courtship in bioluminescent *Photeros* ostracods"

### Supplemental material

Pilot experiment. To estimate how metabolically costly courtship displays were for *Photeros* sp. "EGD," we first needed to determine their baseline metabolism. Individuals of this species are active during the night, but during the day they probably bury themselves in the sediment near seagrass beds, although their daytime location in the wild has never been discovered. Previous anecdotal experience with *Photeros* and other Luxorina suggests they are in a somewhat quiescent state during the day, as they seem less active and less responsive to stimulation. We wanted to test if such differences in activity were driven by circadian rhythms versus exposure to light. Thus, we designed a pilot experiment to assess routine metabolic rate (RMR) at different times (day/night) under different light exposures (ambient light vs foil-wrapped dark).

We measured oxygen consumption as a proxy for metabolic rate, following the methodology explained in the main text. We placed males in 2 mL vessels because we knew, anecdotally, that isolated individuals confined in a small space would not perform courtship displays and were less enticed to swim. We ran trials twice within 24h, recording data once from 11am-5pm (day) and once from 6pm-12am (night). We used two Firesting-O2 4-channel units, each equipped with a temperature probe. In each brick, channels 1 and 2 recorded from chambers exposed to ambient light, while channels 3 and 4 recorded from chambers wrapped in tinfoil. We logged oxygen concentration every 2s using the Pyro Oxygen Logger software (PyroScience, Aachen, Germany). For the data analysis, we used the '*respirometry*' package in R (Birk, 2024) to estimate RMR, accounting for the seawater salinity (33), chamber volume (2 mL), and individual dry mass (0.25 mg; average mass of the males we weighed). We excluded the first hour of recording to minimise effects of animal handling stress on estimations and to allow sufficient mixing of oxygen within the chamber. We did not deduct microbial respiration from the trials because we did not run control chambers as part of the pilot experiment. Lastly, we ran a multiple linear regression to test the effects of time of day, chamber wrapping, and the interaction of these with RMR.

In this pilot experiment, RMR was substantially higher at night than during the day (time of day: $p < 0.001$, df = 12, t = 7.514; Fig. S1). We did not detect a statistically significant effect of chamber wrapping ($p = 0.9583$, df = 12, t = 0.053) or a time-of-day × wrapping interaction ($p = 0.4297$, df = 12, t = 0.817). However, because this pilot experiment had low sample size, the absence of a significant wrapping effect should not be interpreted as strong evidence that darkness has no effect on metabolism. Instead, these results indicate that diel state had a large effect under the conditions tested, while any additional effect of acute chamber darkness was either small relative to the diel effect or not resolvable with the power of this pilot design.

These results suggested that there was a measurable difference in energy expenditure of males between day and night and that RMR was closely tied to activity rates observed for males under field and laboratory conditions; these differences seem to be associated more with time of day than exposure to ambient light. This rough pilot experiment helped inform the overall experimental design because it suggested that periods of inactivity/rest during the night, when individuals are typically more active, do not equate to baseline metabolism. Rather, states akin to torpor during the day, even in the absence of light, greatly reduce energy expenditures and their associated metabolic costs.

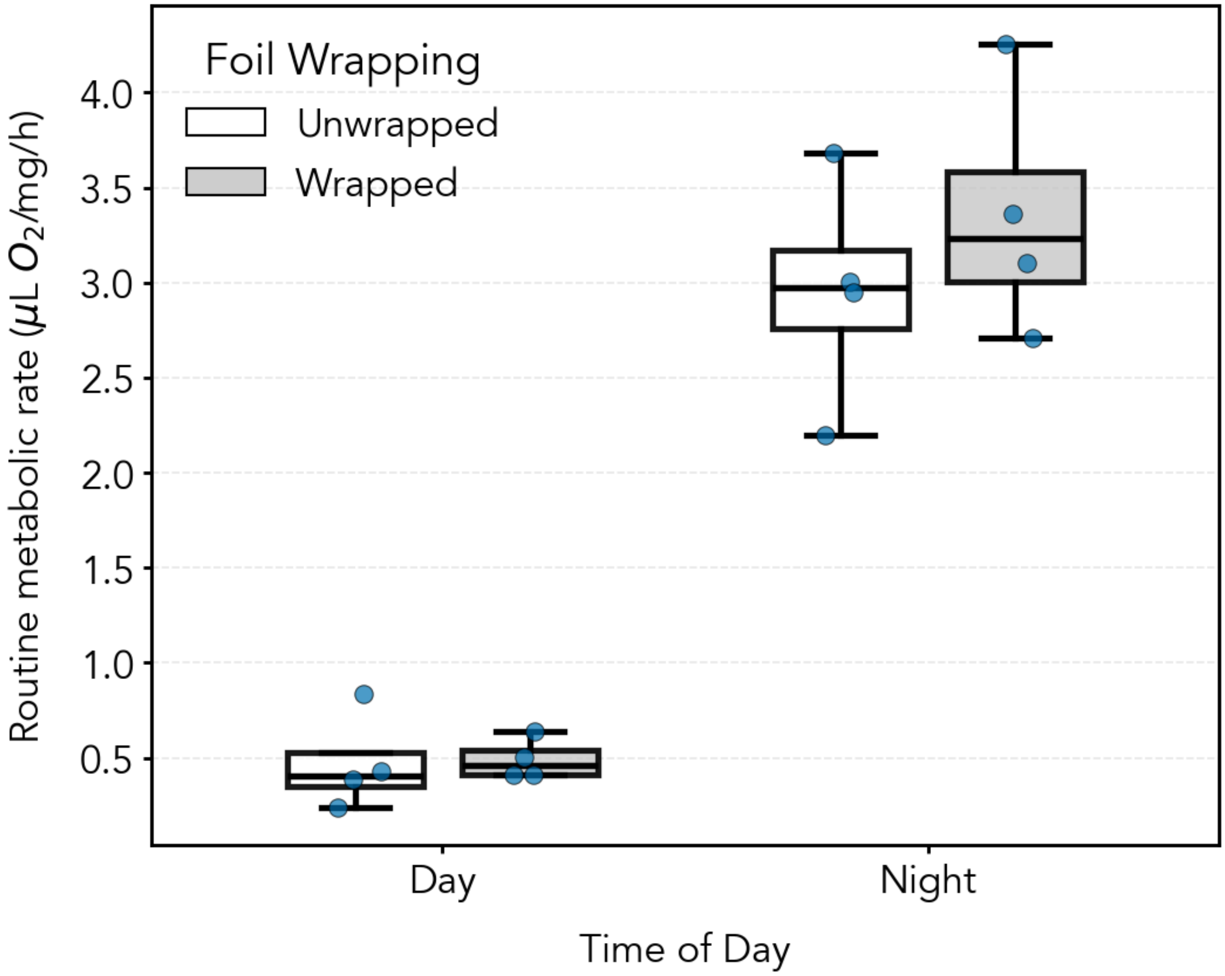


**Fig. S1**. Pilot experiment to measure routine metabolic rate of male ostracods at different times of day and under different aluminium foil-wrapping conditions (i.e., exposures to ambient light). Note these measurements did use a control without animals, and therefore we could not subtract background rates from microbial oxygen consumption. These values then would include any diel differences from microbial consumption, not just oxygen consumption from ostracods.

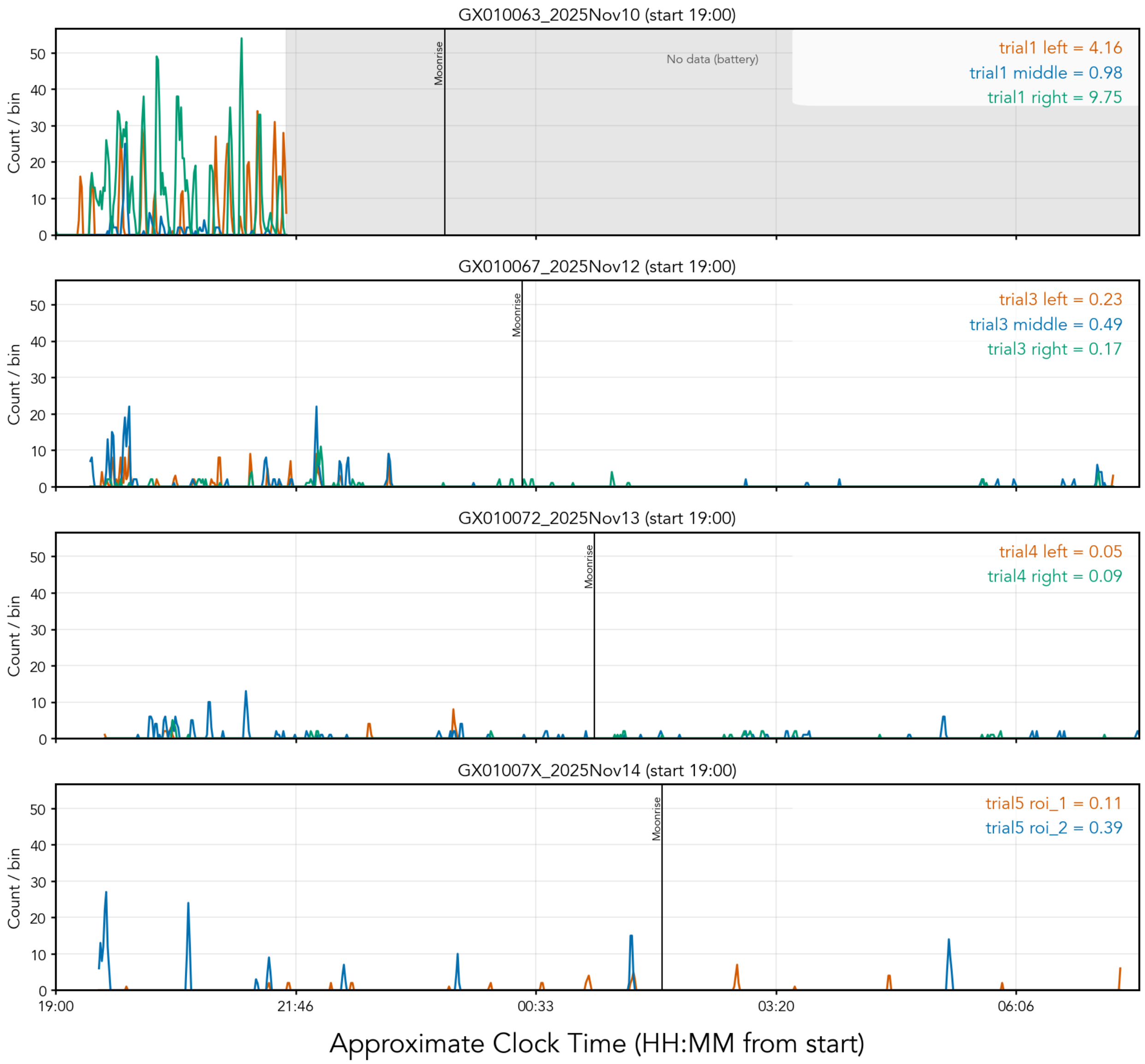


**Figure S2.** Stacked time-series of pulse detections per large vessel as defined in a Region Of Interest (ROI) in video analyses (count per 60 s bin) across four recording nights. Each panel shows one night, with coloured lines indicating vessel positions/ROIs (orange, blue, green), and a vertical black line indicating natural moonrise time (although animals were indoors and covered during experiments). The text in the upper right of each subplot reports the estimate of signal rate for each vessel/ROI, calculated as the mean detections per 60 s bin for that night (for example, `trial1 left = 4.16`). These rate estimates are the key summary values used for downstream activity-metabolism comparisons.

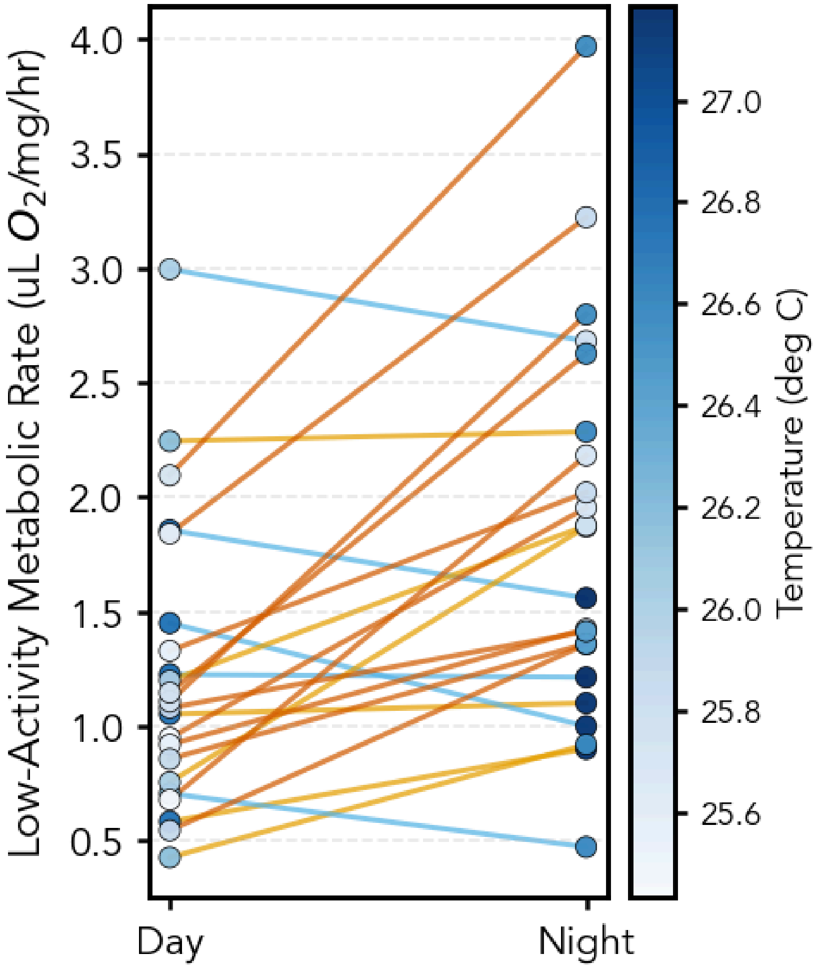


**Figure S3. Temperature did not affect day-night differences of low-activity metabolic rate.** In paired small-vessel measurements, adding temperature to an environment-only random-intercept model did not improve model fit (likelihood-ratio $\chi^2 = 0.413$, df = 1, p = 0.520; temperature estimate = −0.175, p = 0.513; n = 44 measurements from 22 pairs). The same conclusion held when restricted to filtered rows only (p = 0.308), indicating that the day–night difference in low-activity metabolic rate was not better explained by including temperature variation among paired trials. By contrast (see Figure S4), temperature was strongly correlated with signalling rate in large-vessel trials (r = 0.873). Mean temperature was not different between diel pairs, with a mean of 26.03 deg C during day trials and 26.39 deg C during night trials. Because temperature measurements were shared within trial blocks, we tested day-night differences at the trial-pair level (n=4 trial pairs): mean night-day difference=0.37 deg C; paired t-test: t=2.964, p=0.0594; Wilcoxon signed-rank: W=0.000, p=0.1250.These results indicate that temperature does not explain the paired small-vessel diel effect, but that it complicates interpretation of the relationship between signalling activity and oxygen consumption in large vessels.

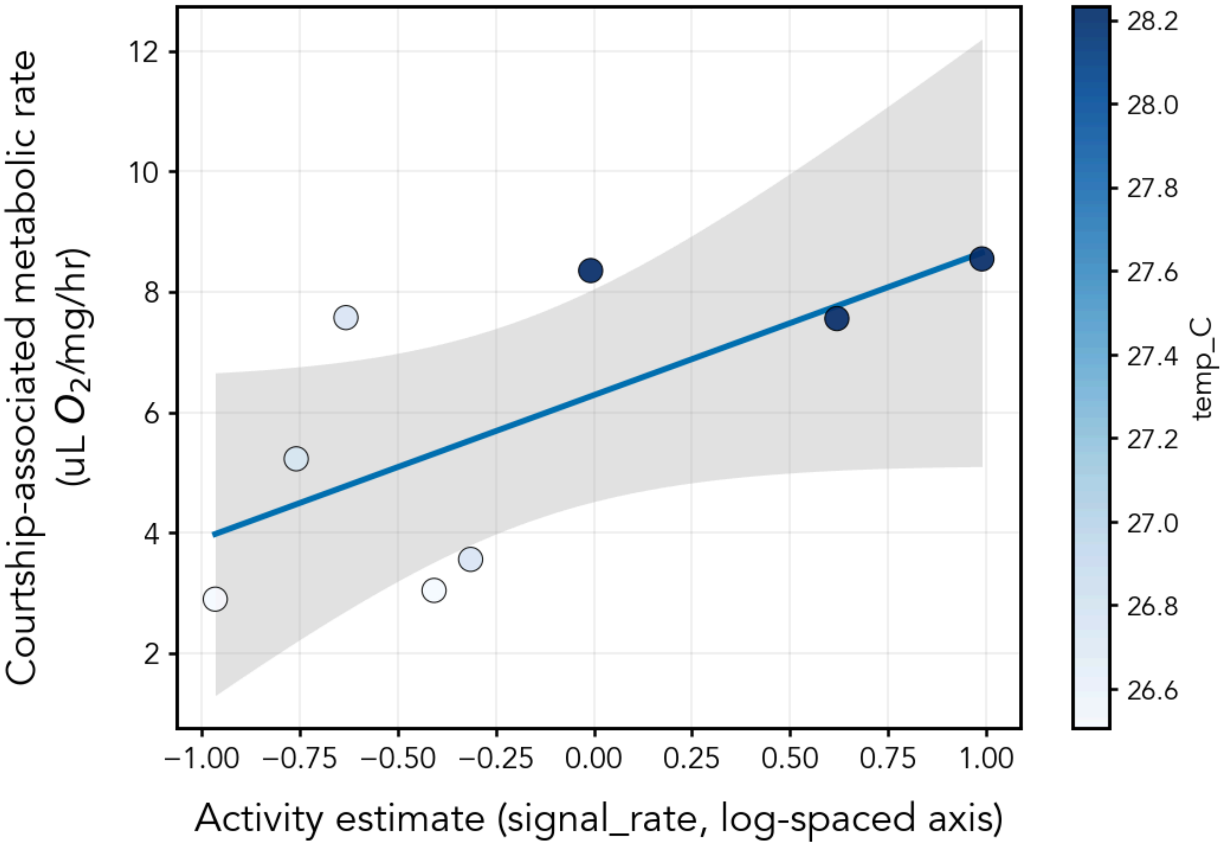


**Figure S4.** Relationship between activity estimate and routine metabolic rate in large-vessel trials while also showing temperature. We found temperature to be strongly correlated with signal rate ($r = 0.873$). In a temperature-adjusted model, this changes the inferred relationship between signal rate and RMR. Using RMR ~ log10(signal_rate) + temp_C, the effect of signal rate was no longer positive or statistically supported ($\beta = -0.838$, $p = 0.661$), while temperature showed a positive but non-significant association with RMR ($\beta = 3.059$, $p = 0.095$; $n = 8$). This suggests that temperature covaries with activity estimates in the large-vessel trials and should be considered when interpreting the apparent activity-RMR relationship. These volatile results again highlight the low sample size and low power when looking for a specific relationship between signalling rate and metabolic rates.

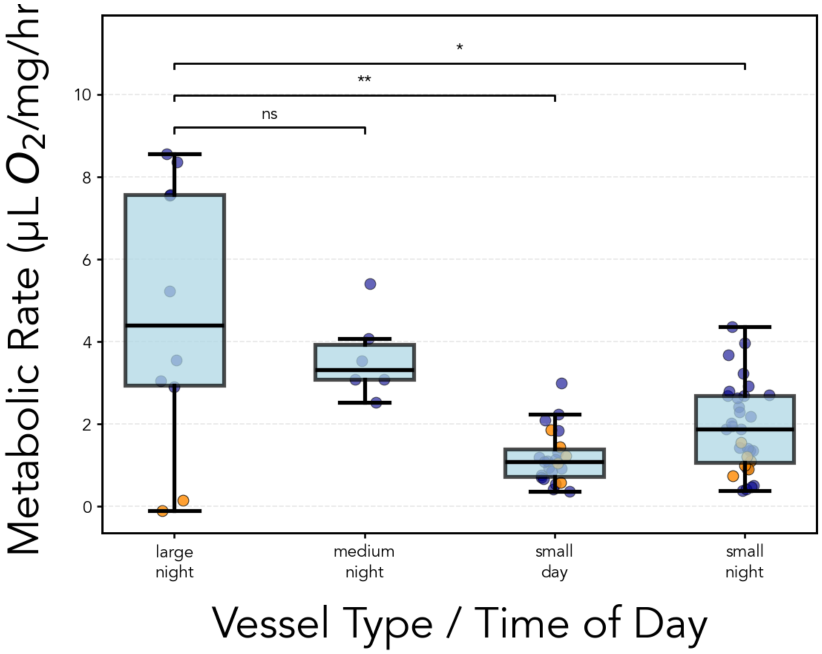


**Figure S5.** Mass-specific metabolic rate (μL O2 mg^-1 h^-1) of *Photeros* across vessel type and time of day, shown for the full dataset including trials excluded from the main analysis. Patterns were consistent with the main figure, with highest rates in large-night trials, intermediate rates in medium-night trials, and lower rates in small-vessel trials, with small-night generally above small-day. Boxplots show medians and interquartile ranges; whiskers show the non-outlier range; points show individual observations. Blue points indicate observations included in the primary analysis, and orange points indicate observations excluded from the primary analysis (for example, trials affected by clogged-filter conditions). Sample sizes in this full dataset were: large-night, n = 10; medium-night, n = 6; small-day, n = 23; small-night, n = 32. Small-vessel data include paired day/night trials as in the main figure plus additional unpaired trials, predominantly at night.

**Table S1.** Temperature-adjusted treatment model for comparisons in main text Figure 2, adding temperature as a co-variate. Model: RMR ~ group + temp_C; reference group = large-night; SE = HC3. Hypothesis test direction: large-night > comparison group; Holm correction applied.

| Comparison | Ncyl | Ncomp | Mean cyl | Mean comp | Adjusted diff. | Adjusted one-sided p | Holm-adjusted p | Signif? |
|---|---|---|---|---|---|---|---|---|
| Large vs medium | 8 | 6 | 5.844 | 3.615 | 2.344 | 0.007 | 0.007 | Yes |

| | | | | | | | | |
|---|---|---|---|---|---|---|---|---|
| Large vs. small (day) | 8 | 18 | 5.844 | 1.181 | 3.566 | <0.001 | <0.001 | Yes |
| Large vs. small (night) | 8 | 26 | 5.844 | 2.098 | 3.297 | <0.001 | <0.001 | Yes |

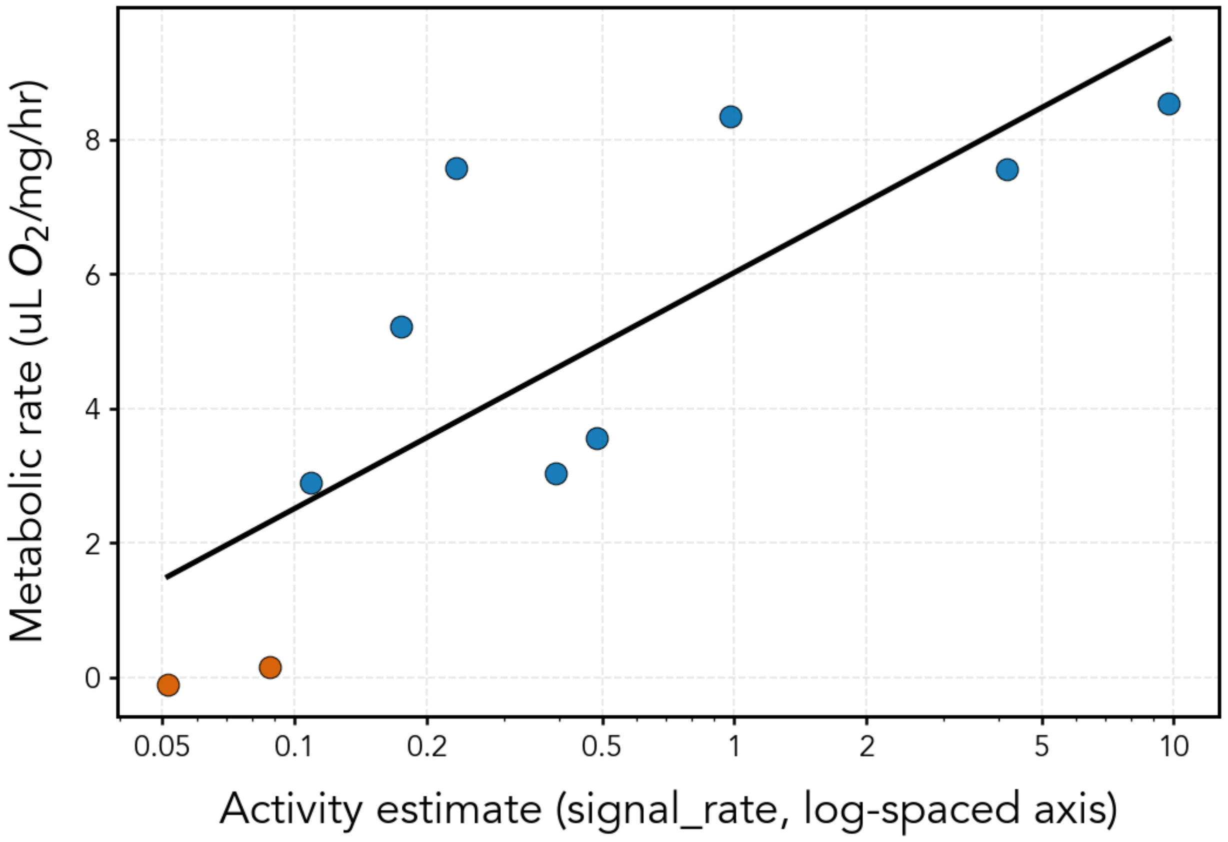


**Figure S6.** Relationship between activity estimate and metabolic rate in large-vessel trials when all data are included, including respiration-rate values excluded from the main analysis. Blue points indicate trials retained in the main text analysis (filtered=True), and orange points indicate trials excluded from the main text (filtered=False; including negative-valued excluded observations). Metabolic rate (RMR) is plotted against activity estimate (signal_rate; log-scaled x-axis), and the solid line shows the least-squares fit to log transformed activity estimates across all included points in this supplementary analysis. The relationship is positive (slope = 3.51), with moderate-to-strong explained variance ($R^2$ = 0.62) and significant statistical support ((p=0.0066, n=10)).

<u>Updating the signalling-cost comparison.</u> We updated the Stoddard and Salazar (2011) signalling-cost dataset to place *Photeros* sp. EGD courtship metabolism in a broader

comparative context. We used Deep Research (ChatGPT) to identify post-2011 studies measuring metabolic rates during animal signalling and during a resting, silent, or low-activity baseline, and then audited candidate papers using original publications, supplementary files, and public repositories. Search terms combined “oxygen consumption,” “metabolic rate,” “$VO_2$,” “$\dot{V}O_2$,” “$VCO_2$,” “$CO_2$ production,” and “respirometry” with signalling terms such as “calling,” “song,” “whistle,” “click,” “echolocation,” “hissing,” “electric organ discharge,” “stridulation,” and “vibrational signalling.”

Studies were retained only when signalling and baseline metabolic rates could be expressed as mass-specific oxygen consumption in the same units as the legacy dataset, ml $O_2$ $h^{-1}$ $g^{-1}$. Values reported as ml $O_2$ $min^{-1}$ $kg^{-1}$ were multiplied by 0.06. Whole-animal rates were divided by body mass when body mass was reported for the same animals. One $CO_2$-based study was converted to $O_2$ using an assumed respiratory quotient of 0.85 and flagged as converted. Studies were excluded from the strict update when they reported only relative changes, lacked a resting or signalling metabolic rate, measured signalling and resting in non-comparable contexts, or reported only recovery-inclusive energetic costs that could not be converted to a signalling-period rate.

The Deep Research search identified three rows that were immediately compatible with the previously published meta-analysis: *Plangia graminea* calling (Doubell et al., 2017), bottlenose dolphin whistling (Pedersen et al., 2020), and *Thyroptera tricolor* social calling (Chaverri et al., 2021). Human-assisted extraction added four additional rows: bottlenose dolphin communicative sounds from (Noren et al., 2013), bottlenose dolphin echolocation clicks from (Noren et al., 2017), *Aphrodes makarovi* vibrational advertisement calls from (Kuhelj et al., 2015), and *Vipera ammodytes* defensive hissing from (Van Zele et al., 2024). The dolphin echolocation row was calculated from the published supplementary spreadsheet as submerged clicking metabolism divided by submerged silent metabolism. The viper hissing row was calculated from the Zenodo respirometry dataset using paired individuals with both control and hissing measurements. The leafhopper row used directly reported mass-specific $O_2$ values for resting and advertisement-call production.

We retained the expanded dataset as a descriptive comparison rather than an inferential meta-analysis. The added studies differ in signal modality, respirometry method, baseline definition, and whether signalling rates represent instantaneous signal production, trial-averaged behaviour, or mixed-call trials. Extraction notes and flags were therefore preserved in the archived CSV files.

**Table S2.** Unified data table. Trial=unique trial number; brick=the recording box; channel=one of four channel numbers on each brick; n=number of animals in the vessel; total_mass=the measured mass of the animals in the trial; temp_C=temperature measured from probe; vessel=size of the container (cylinder = large); date=date of trial; envi=time of the trial; either day or night; filtered=whether water was filtered or not; signal_rate=estimate of signalling activity from video analysis; pair=partner of a paired trial

| trial | brick | channel | n | total_mass | RMR | temp_C | vessel | date | envi | filtered | signal_rate | pair |
|---|---|---|---|---|---|---|---|---|---|---|---|---|
| trial1 | box2 | Ch2 | 1 | 0.2263 | 2.708969368 | 27.948 | small | 10Nov2025 | night | TRUE | | |
| trial1 | box2 | Ch3 | 1 | 0.2099 | 3.669334636 | 27.948 | small | 10Nov2025 | night | TRUE | | |
| trial1 | box2 | Ch4 | 1 | 0.2703 | 2.927729211 | 27.948 | small | 10Nov2025 | night | TRUE | | |
| trial1 | box3 | Ch2 | 20 | 7.6 | 8.546659523 | 28.233 | cylinder | 10Nov2025 | night | TRUE | 9.745341615 | |
| trial1 | box3 | Ch3 | 20 | 6.5 | 8.352764258 | 28.233 | cylinder | 10Nov2025 | night | TRUE | 0.9813664596 | |
| trial1 | box3 | Ch4 | 20 | 5.5 | 7.555755104 | 28.233 | cylinder | 10Nov2025 | night | TRUE | 4.161490683 | |
| trial1 | newbox | Ch2 | 1 | 0.2513 | 2.689698807 | 27.921 | small | 10Nov2025 | night | TRUE | | |
| trial1 | newbox | Ch3 | 1 | 0.297 | 2.420259881 | 27.921 | small | 10Nov2025 | night | TRUE | | |
| trial1 | newbox | Ch4 | 1 | 0.1811 | 4.356230825 | 27.921 | small | 10Nov2025 | night | TRUE | | |
| trial2 | newbox | Ch2 | 5 | 1.1 | 3.09154531 | 26.846 | medium | 11Nov2025 | night | TRUE | | |
| trial2 | newbox | Ch3 | 5 | 0.9 | 5.398751174 | 26.846 | medium | 11Nov2025 | night | TRUE | | |
| trial2 | newbox | Ch4 | 5 | 1.1 | 3.530252687 | 26.846 | medium | 11Nov2025 | night | TRUE | | |
| trial3 | box2 | Ch2 | 20 | 4.9 | 5.227565289 | 26.75 | cylinder | 12Nov2025 | night | TRUE | 0.1744022504 | |
| trial3 | box2 | Ch3 | 20 | 4.7 | 3.558230156 | 26.75 | cylinder | 12Nov2025 | night | TRUE | 0.4852320675 | |
| trial3 | box2 | Ch4 | 20 | 4.3 | 7.572331593 | 26.75 | cylinder | 12Nov2025 | night | TRUE | 0.2334739803 | |
| trial3 | box3 | Ch2 | 5 | 1.1 | 3.076208738 | 27.948 | medium | 12Nov2025 | night | TRUE | | |
| trial3 | box3 | Ch3 | 5 | 1 | 2.521120116 | 27.948 | medium | 12Nov2025 | night | TRUE | | |
| trial3 | box3 | Ch4 | 5 | 0.9 | 4.07079645 | 27.948 | medium | 12Nov2025 | night | TRUE | | |
| trial3 | newbox | Ch2 | | 0.2168 | 0.3842837414 | 26.813 | small | 12Nov2025 | night | TRUE | | |
| trial3 | newbox | Ch3 | | 0.1989 | 0.5170944169 | 26.813 | small | 12Nov2025 | night | TRUE | | |
| trial3 | newbox | Ch4 | | 0.2552 | 0.4283317982 | 26.813 | small | 12Nov2025 | night | TRUE | | |
| trial4 | box2 | Ch2 | | 0.2187 | 1.101950466 | 27.142 | small | 13Nov2025 | night | FALSE | | trial4.5 |
| trial4 | box2 | Ch3 | | 0.3741 | 0.9014938545 | 27.142 | small | 13Nov2025 | night | FALSE | | trial4.5 |
| trial4 | box2 | Ch4 | | 0.2074 | 1.000535975 | 27.142 | small | 13Nov2025 | night | FALSE | | trial4.5 |
| trial4 | box3 | Ch2 | | 0.2946 | 0.7405659426 | 27.185 | small | 13Nov2025 | night | FALSE | | trial4.5 |
| trial4 | box3 | Ch3 | | 0.2209 | 1.214074578 | 27.185 | small | 13Nov2025 | night | FALSE | | trial4.5 |
| trial4 | box3 | Ch4 | | 0.2518 | 1.56000791 | 27.185 | small | 13Nov2025 | night | FALSE | | trial4.5 |
| trial4 | newbox | Ch2 | 20 | 4.5 | 0.1580963749 | 27.174 | cylinder | 13Nov2025 | night | FALSE | 0.0876216968 | trial4.5 |
| trial4 | newbox | Ch4 | 20 | 4.9 | -0.1030399276 | 27.174 | cylinder | 13Nov2025 | night | FALSE | 0.05146036161 | trial4.5 |
| trial4.5 | box2 | Ch2 | | 0.2187 | 1.053823758 | 26.728 | small | 14Nov2025 | day | FALSE | | trial4 |

| | | | | | | | | | | | | |
|---|---|---|---|---|---|---|---|---|---|---|---|---|
| trial4.5 | box2 | Ch3 | | 0.3741 | 0.5844811658 | 26.728 | small | 14Nov2025 | day | FALSE | | trial4 |
| trial4.5 | box2 | Ch4 | | 0.2074 | 1.450557575 | 26.728 | small | 14Nov2025 | day | FALSE | | trial4 |
| trial4.5 | box3 | Ch3 | | 0.2209 | 1.226848728 | 26.774 | small | 14Nov2025 | day | FALSE | | trial4 |
| trial4.5 | box3 | Ch4 | | 0.2518 | 1.85686262 | 26.774 | small | 14Nov2025 | day | FALSE | | trial4 |
| trial5 | box2 | Ch2 | 20 | 3.3 | 3.042832094 | 26.506 | cylinder | 14Nov2025 | night | TRUE | 0.3915492958 | trial5.5 |
| trial5 | box2 | Ch4 | 20 | 3.6 | 2.897411334 | 26.506 | cylinder | 14Nov2025 | night | TRUE | 0.1084507042 | trial5.5 |
| trial5 | box3 | Ch2 | | 0.1893 | 2.285837383 | 26.568 | small | 14Nov2025 | night | TRUE | | trial5.5 |
| trial5 | box3 | Ch3 | | 0.3351 | 0.9208535111 | 26.568 | small | 14Nov2025 | night | TRUE | | trial5.5 |
| trial5 | box3 | Ch4 | | 0.3627 | 0.4736240867 | 26.568 | small | 14Nov2025 | night | TRUE | | trial5.5 |
| trial5 | newbox | Ch2 | | 0.2234 | 1.871534338 | 25.773 | small | 14Nov2025 | night | TRUE | | trial5.5 |
| trial5 | newbox | Ch3 | | 0.2704 | 1.876437076 | 25.773 | small | 14Nov2025 | night | TRUE | | trial5.5 |
| trial5 | newbox | Ch4 | | 0.2277 | 2.681803129 | 25.773 | small | 14Nov2025 | night | TRUE | | trial5.5 |
| trial5.5 | newbox | Ch2 | | 0.2234 | 0.7561932576 | 26.004 | small | 15Nov2025 | day | TRUE | | trial5 |
| trial5.5 | newbox | Ch3 | | 0.2704 | 1.199553463 | 26.004 | small | 15Nov2025 | day | TRUE | | trial5 |
| trial5.5 | newbox | Ch4 | | 0.2277 | 2.995435266 | 26.004 | small | 15Nov2025 | day | TRUE | | trial5 |
| trial5.5 | box3 | Ch2 | | 0.1893 | 2.245259005 | 26.159 | small | 15Nov2025 | day | TRUE | | trial5 |
| trial5.5 | box3 | Ch3 | | 0.3351 | 0.4278883728 | 26.159 | small | 15Nov2025 | day | TRUE | | trial5 |
| trial5.5 | box3 | Ch4 | | 0.3627 | 0.706451259 | 26.159 | small | 15Nov2025 | day | TRUE | | trial5 |
| trial6 | box3 | Ch2 | | 0.2401 | 0.9498575116 | 25.433 | small | 16Nov2025 | day | TRUE | | trial6.5 |
| trial6 | box3 | Ch3 | | 0.2579 | 0.3753363284 | 25.433 | small | 16Nov2025 | day | TRUE | | trial6.5 |
| trial6 | box3 | Ch4 | | 0.2079 | 0.6794722856 | 25.433 | small | 16Nov2025 | day | TRUE | | trial6.5 |
| trial6 | newbox | Ch2 | | 0.193 | 1.841349067 | 25.551 | small | 16Nov2025 | day | TRUE | | trial6.5 |
| trial6 | newbox | Ch3 | | 0.2411 | 0.9202794488 | 25.551 | small | 16Nov2025 | day | TRUE | | trial6.5 |
| trial6 | newbox | Ch4 | | 0.2281 | 1.329196664 | 25.551 | small | 16Nov2025 | day | TRUE | | trial6.5 |
| trial6.5 | box3 | Ch2 | | 0.2401 | 1.954137166 | 25.716 | small | 16Nov2025 | night | TRUE | | trial6 |
| trial6.5 | box3 | Ch4 | | 0.2079 | 2.182509445 | 25.716 | small | 16Nov2025 | night | TRUE | | trial6 |
| trial6.5 | newbox | Ch2 | | 0.193 | 3.222056421 | 25.838 | small | 16Nov2025 | night | TRUE | | trial6 |
| trial6.5 | newbox | Ch3 | | 0.2411 | 1.424906572 | 25.838 | small | 16Nov2025 | night | TRUE | | trial6 |
| trial6.5 | newbox | Ch4 | | 0.2281 | 2.021198508 | 25.838 | small | 16Nov2025 | night | TRUE | | trial6 |
| trial7 | newbox | Ch2 | | 0.2279 | 2.096022083 | 25.734 | small | 17Nov2025 | day | TRUE | | trial7.5 |
| trial7 | newbox | Ch3 | | 0.2633 | 1.108404679 | 25.734 | small | 17Nov2025 | day | TRUE | | trial7.5 |
| trial7 | newbox | Ch4 | | 0.2362 | 1.147038409 | 25.734 | small | 17Nov2025 | day | TRUE | | trial7.5 |
| trial7 | box3 | Ch2 | | 0.2644 | 0.5460283033 | 25.873 | small | 17Nov2025 | day | TRUE | | trial7.5 |
| trial7 | box3 | Ch3 | | 0.2535 | 0.8577576504 | 25.873 | small | 17Nov2025 | day | TRUE | | trial7.5 |
| trial7 | box3 | Ch4 | | 0.249 | 1.08048547 | 25.873 | small | 17Nov2025 | day | TRUE | | trial7.5 |
| trial7.5 | newbox | Ch2 | | 0.2279 | 3.967866999 | 26.549 | small | 17Nov2025 | night | TRUE | | trial7 |
| trial7.5 | newbox | Ch3 | | 0.2633 | 2.798130052 | 26.549 | small | 17Nov2025 | night | TRUE | | trial7 |
| trial7.5 | newbox | Ch4 | | 0.2362 | 2.626962253 | 26.549 | small | 17Nov2025 | night | TRUE | | trial7 |
| trial7.5 | box3 | Ch2 | | 0.2644 | 1.358387376 | 26.417 | small | 17Nov2025 | night | TRUE | | trial7 |
| trial7.5 | box3 | Ch3 | | 0.2535 | 1.360359664 | 26.417 | small | 17Nov2025 | night | TRUE | | trial7 |
| trial7.5 | box3 | Ch4 | | 0.249 | 1.414542953 | 26.417 | small | 17Nov2025 | night | TRUE | | trial7 |
